# Combined representation of visual features in the scene-selective cortex

**DOI:** 10.1101/2023.07.24.550280

**Authors:** Jisu Kang, Soojin Park

## Abstract

Visual features of separable dimensions like color and shape conjoin to represent an integrated entity. We investigated how visual features bind to form a complex visual scene. Specifically, we focused on features important for visually guided navigation: direction and distance. Previously, separate works have shown that directions and distances of navigable paths are coded in the occipital place area (OPA). Using functional magnetic resonance imaging (fMRI), we tested how separate features are concurrently represented in the OPA. Participants saw eight different types of scenes, in which four of them had one path and the other four had two paths. In single-path scenes, path direction was either to the left or to the right. In double-path scenes, both directions were present. Each path contained a glass wall located either near or far, changing the navigational distance. To test how the OPA represents paths in terms of direction and distance features, we took three approaches. First, the independent-features approach examined whether the OPA codes directions and distances independently in single-path scenes. Second, the integrated-features approach explored how directions and distances are integrated into path units, as compared to pooled features, using double-path scenes. Finally, the integrated-paths approach asked how separate paths are combined into a scene. Using multi-voxel pattern similarity analysis, we found that the OPA’s representations of single-path scenes were similar to other single-path scenes of either the same direction or the same distance. Representations of double-path scenes were similar to the combination of two constituent single-paths, as a combined unit of direction and distance rather than pooled representation of all features. These results show that the OPA combines the two features to form path units, which are then used to build multiple-path scenes. Altogether, these results suggest that visually guided navigation may be supported by the OPA that automatically and efficiently combines multiple features relevant for navigation and represent a *navigation file*.

## INTRODUCTION

We instantly see the world and objects in it as complete entities, without having to first analyze the features that compose them. For instance, it is not necessary in our daily lives to inspect in detail the round-shape, red-color, fist-size, and the waxy-texture of an object to conclude that it looks like an apple. We do not have to identify each of the buildings first to then perceive a coherent view of the city. In fact, this ability of ours is supported by the visual system that rapidly puts together small pieces of information to perceive complete entities (Humphreys & Riddoch, 2012). Visual inputs are first processed in the early visual cortex (EVC) of the brain as simple lower-level features like color, size, and orientation (Hubel and Wiesel, 1959). These features of separate dimensions are later integrated as the information is passed along the visual pathway to form more complex features (Felleman & Van Essen, 1991; Grill-Spector, Kushnir, Itzchak, & Malach, 1998; Kravitz, Saleem, Baker, & Mishkin, 2011).

Furthermore, the visual system can combine small object parts to represent a larger complex object. For instance, it has been demonstrated that the response of the fusiform gyrus to a whole person is best modeled by the linear combination of independent face and body parts with equal weights (Kaiser et al., 2014). With a similar logic, Erez, Kendall, & Barense (2016) showed that the lateral occipital complex (LOC) extending into the perirhinal cortex (PRC) has conjoined coding of smaller parts of an object in representing a larger complex object. Here, the experimenters used artificially created objects that could have any of the three smaller parts attached to them (parts A, B, and C). Then they combined the representations of an object with part A and an object with parts BC (A + BC) to compare with the combination of objects with part B and parts AC (B + AC), and the combination of objects with part C and parts AB (C + AB). The parts included in all combinations were equal, but the specific conjunctions of parts differed. The authors found representations for the precise conjunctions of the parts of an object in the LOC and the PRC.

Coding of features is not only important for object recognition, but it is also critical for a complete visual scene recognition. The importance of junctions and curvatures was revealed by comparing the error patterns of scene categorization by humans and computers (Walther & Shen, 2014). Moreover, human behavioral performance on categorization decreased when depriving of long contours from line drawings of scenes (Walther, Chai, Caddigan, Beck, & Fei-Fei, 2011). Scene-selective cortices also code for features that reside in visual scenes. Using scene images and simple geometrical images, Nasr and Tootell (2012) found a higher activity level in the scene-selective parahippocampal place area (PPA) for cardinal orientations, which are predominant features of scenes, compared to oblique orientations. Geometrical features are also coded in the PPA, shown by its stronger response to an empty room than the same room with its walls, floor, and ceiling fractured and rearranged (Epstein & Kanwisher, 1998).

Evidence suggests that the occipital place area (OPA), another scene selective region, also represents features that are especially relevant for navigation. First, the OPA was found to be sensitive to identical images that were mirror-reversed, which critically changed the direction of paths while other features stayed the same (Dilks, Julian, Kubilius, Spelke, & Kanwisher, 2011). Directions and distances of spatial boundaries are examples of independent visual feature dimensions that reside in scenes and are highly relevant for visually guided navigation. With respect to direction features, Bonner and Epstein (2017) showed that the OPA codes navigational affordances in visual scenes, which referred to the navigational trajectories and their angular directions. They were able to reconstruct path trajectories from the response patterns of the OPA. More evidence revealed that the OPA strictly distinguishes between navigable distances, which is the distance from self to a spatial boundary structure that limits locomotion like glass walls, and not transparent curtains (Park & Park, 2020).

While the mechanisms behind the human ability to represent individual features and perceive integrated objects have been studied extensively, the mechanism underlying the visual perception of a scene as a complete set of features, especially those related to navigation, is less investigated. The ability to bind navigationally relevant features is especially important for planning and executing of motion. Correct integration of the orientations and lengths of paths is necessary to determine where and how far one can move. If the visual system fails to accurately combine directions and distances of paths, then it would be difficult to calculate possible navigational trajectories which requires these features simultaneously. Here, we aim to investigate how the two navigationally relevant features, directions and distances, are concurrently represented in the OPA. We used artificially created scene images with three conditions. There were two levels in each condition, creating a total of eight stimulus conditions (number of paths: single-path and double-path, directions: left and right, and distances: near and far). fMRI multi-voxel patterns were collected from the OPA while the participants viewed the stimuli. Neural similarity analysis was used to investigate how these scene regions represent visual features of path directions and distances in unambiguous single-path scenes and complex double-path scenes. We took three different approaches, all using the neural similarity method. First, in the *independent-features* approach, we test whether direction features and distance features in our stimuli are coded by the OPA with no preference for one feature over another. An additional analysis compares the response patterns to a low-level image similarity model, to account for the effect of low-level features. Then in the *integrated-features* approach, we investigate whether the OPA represents an integrated form of direction and distance features as a path. In the *integrated-paths* approach, we further inspect whether the coding of multiple paths within a scene combine to represent a relatively complex scene, similar to how the coding of parts of an object conjoin to represent a larger complex object (Erez et al., 2016).

To test if the conjunctive coding of features in scenes is unique to the OPA, we compared the OPA’s response in all of our analyses with the responses of the PPA, another scene-selective region (Epstein & Kanwisher, 1998). The PPA is widely known to represent categories of scenes than its navigational properties. For example, the response of the PPA is stronger when performing a scene categorization task than a navigation task using the same images (Persichetti & Dilks, 2018). It was also found to be sensitive to scene layout, defined by open and closed spatial boundaries, which is a critical component for scene categorization (Park, Brady, Greene, & Oliva, 2011). We selected the PPA as a control area since it is similar to the OPA in a way that it responds strongly to scenes in general but engages in a function other than navigation.

### Independent Representations for Direction and Distance: Independent-features approach

In the *independent-features* approach, we investigated how two features of different dimensions, path direction and distance, are independently represented in the OPA. We assessed whether the multi-voxel pattern similarities are higher for pairs of single-path scenes that share the direction compared to those that do not share the direction and the distance. To do this, we compared the average of multi-voxel pattern similarities of stimulus condition pairs that share only the direction against the average similarities of pairs that do not overlap in both direction and distance (Figure 1). We also examined whether the multi-voxel pattern similarities are higher for single-path scene pairs that share the distance compared to those that do not share any feature. We then compared the average similarity of stimulus condition pairs that share only the direction against the average similarity of pairs that share only the distance.

**Figure 1.**
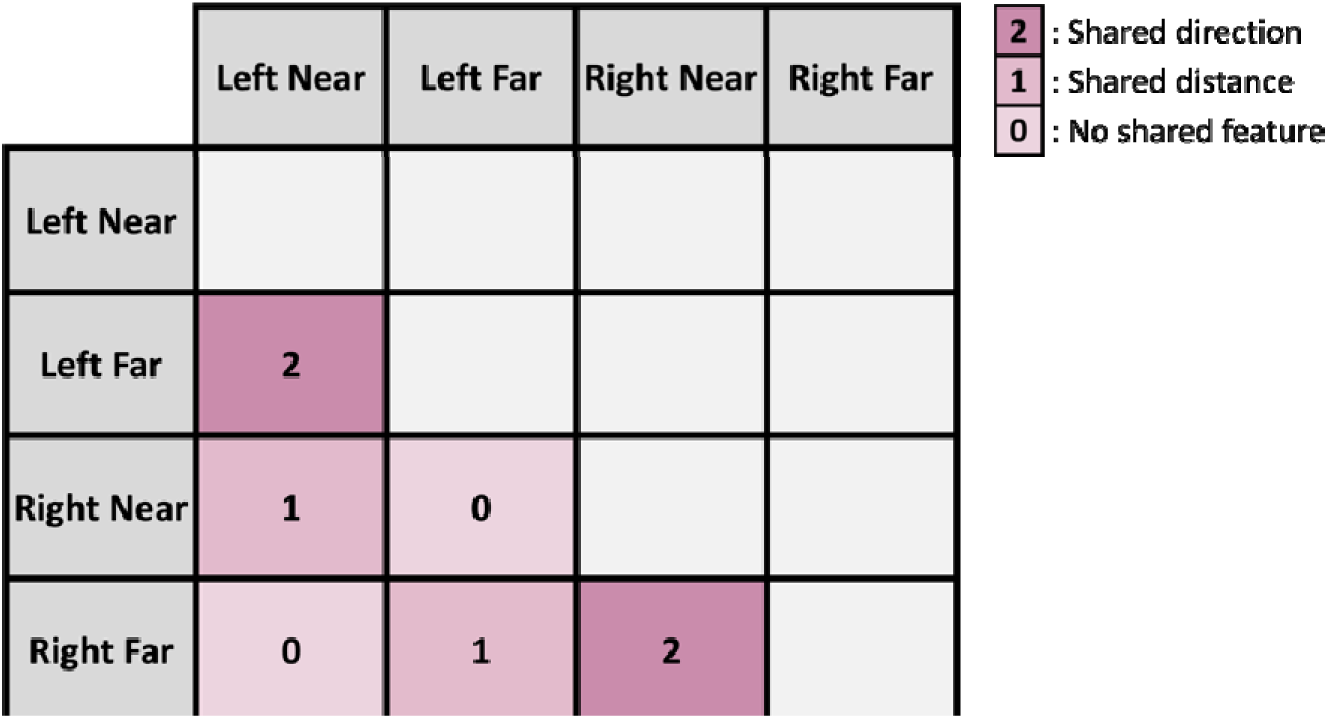
*Independent-features* approach contrast. Contrast conditions assessing the level of voxel-wise neural similarity between single-path scene pairs that share the direction but not the distance (cells marked with 2), and pairs that share the distance but not the direction (cells marked with 1) to pairs that do not share any of the features (cells marked with 0).

The *shared direction* condition is composed of voxel-wise neural similarities of single-path scene pairs that have the same direction but different distances (cells marked with 2 in Figure 1), and the *shared distance* condition includes neural similarities of single-path scene pairs that have the same distance but different directions (cells marked with 1 in Figure 1). The *no shared feature* condition consists of neural similarities of single-path scene pairs that have different directions and different distances (cells marked with 0 in Figure 1).

Figure 2 depicts an example for the three conditions: a *left near* path shares its direction with a *left far* path, its distance with a *right near* path, and it does not share any feature with the *right far* path. If the OPA represents path direction and distance, we hypothesize that the neural similarities of single-path scene pairs in the *no shared feature* condition will be lower compared to those in the *shared direction* condition and compared to the *shared distance* condition.

**Figure 2.**
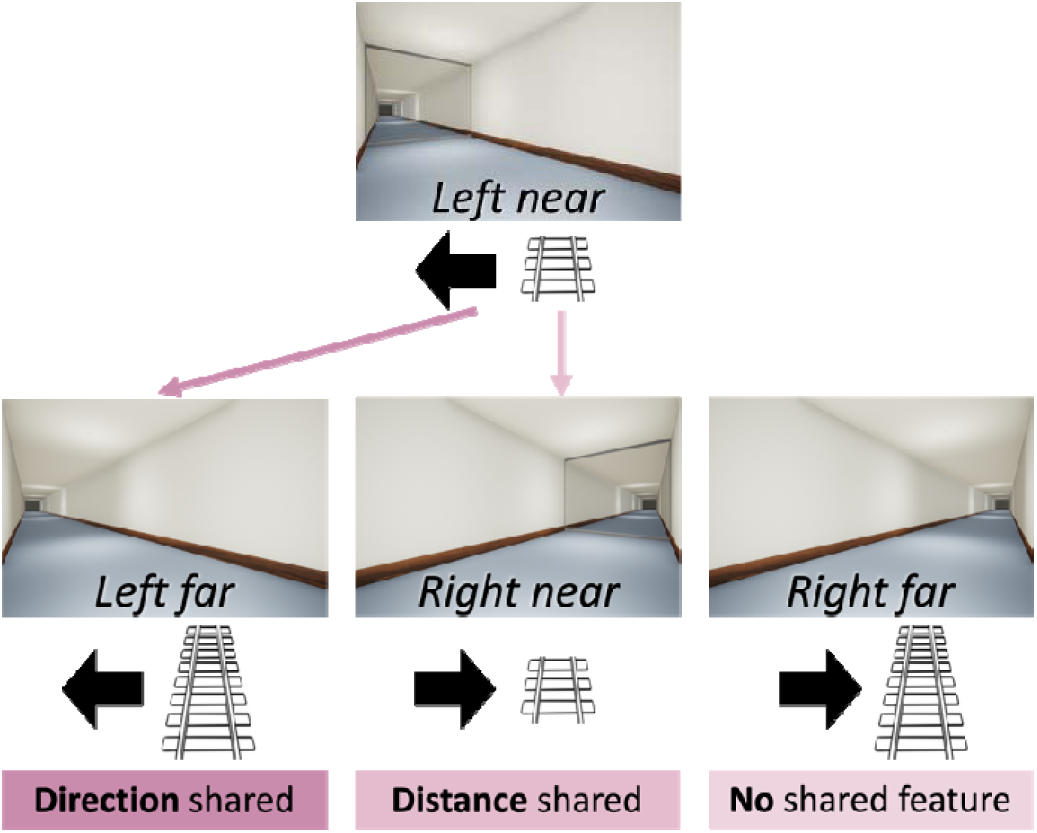
An illustration of an example for the *shared direction*, *shared distance*, and *no shared feature* conditions. The left and right arrows represent left and right path directions. The short and long rails represent near and far distances.

If the OPA has significantly higher neural similarities for both the *shared direction* condition and the *shared distance* condition than the *no shared feature* condition and shows a significant difference between the *shared direction* and the *shared distance* conditions, then it could be interpreted that the OPA’s response is based heavily on one of the two features in representing a path. If the OPA shows a difference between the *no shared feature* condition and the other two conditions but shows no significant difference between the *shared direction* and the *shared distance* conditions, then there is no evidence that the OPA is biased towards representing one feature over the other. We performed a 2 3 ANOVA to confirm differential response patterns between the OPA and the PPA. We then test for differences among the three conditions particularly in the OPA, and significance levels of for each contrast were corrected using false discovery rate (FDR).

### Combined Representation of Directions and Distances: Integrated-features approach

In the *integrated-features* approach, we investigated how directions and distances of multiple paths within a visual scene are concurrently represented in the OPA. We assessed the integration of navigational distances onto path directions in double-path scenes, which would then form a path unit (a combined form of direction and distance). For example, features in a *left-near right-far* double-path can be decomposed into individual features of *left, right, near,* and *far*. The direction features and the distance features could be represented in this decomposed state in the OPA, or they could be combined into two path units. In this case, the features can correctly combine to form a {*left near*} unit and a {*right far*} unit, but they can also incorrectly combine to form a {*left far*} unit and a {*right near*} unit (Figure 3). The OPA’s response patterns for a *left-near right-far* scene may be indistinguishable from the response for a *left-far right-near* scene if features are in a pooled state rather than an integrated state.

**Figure 3.**
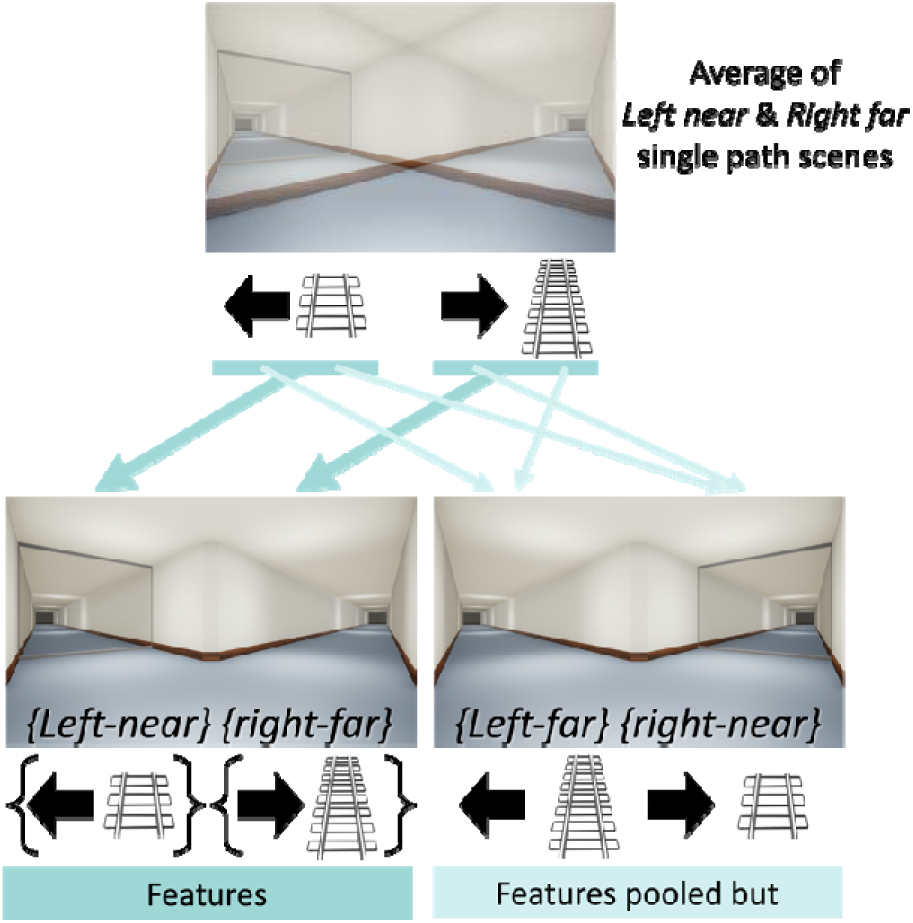
An illustration of an example for the *correctly integrated* condition and the *incorrectly integrated* condition. Brackets represent the conjunction of features.

To test how multiple directions and distances are concurrently coded in the OPA, we conducted an analysis using voxel-based linear combination. Specifically, we synthesized the average representations of two single-path stimulus conditions with different directions and different distances to compare with the representations of double-path scenes (Figure 4a; Baldassano, Beck, & Fei-Fei, 2017; MacEvoy & Epstein, 2009). To do this, we first added the two multi-voxel activity patterns of the two single-path scenes and divided it by two, which resulted in the average representation for each single-path combination. There were two single-path combinations in total (*left near* & *right far* combination and *left far* & *right near* combination). We calculated the correlations between these synthesized average representations and the representations of two double-path scenes with different distances on each path (Figure 4b).

**Figure 4.**
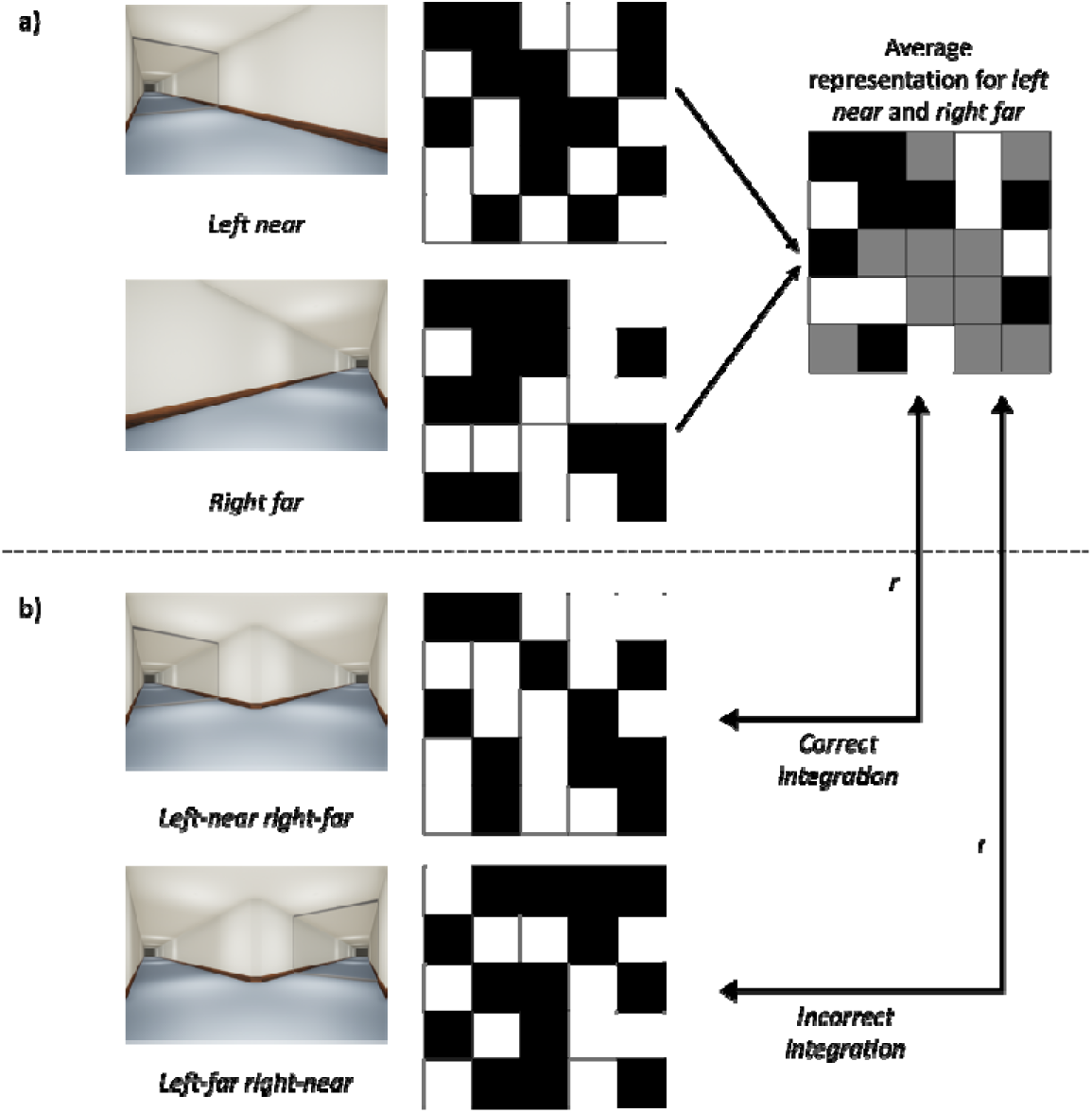
(a) A schematic visualization of an example of the linear combination (average) of the multi-voxel patterns of two single-path scenes of different directions and different distances. (b) Examples of neural similarities between an average representation of two single-path scenes and a double path scene. One example from the *correct integration* condition and another example from the *incorrect integration* condition are depicted.

We then contrasted the neural similarities of synthesized representations and the representations for double-path scenes that share a path unit, as a combined form of direction and distance, with the neural similarities for pairs that do not share any path unit (Figure 5). We focused here on scenes that have different distances on each path to test specifically for the possibility of incorrect combination of features. The *correct integration* condition (cells marked with 1 in Figure 5) consists of neural similarities of scene pairs that share a whole path unit (distance *and* direction). For example, a *left-near right-far* double-path scene shares a whole path unit with the representation synthesized using a *left near* single-path scene and a *right-far* single path scene. The *incorrect integration* condition (cells marked with 0 in Figure 5) includes scene pairs not included in the *correct integration* condition.

**Figure 5.**
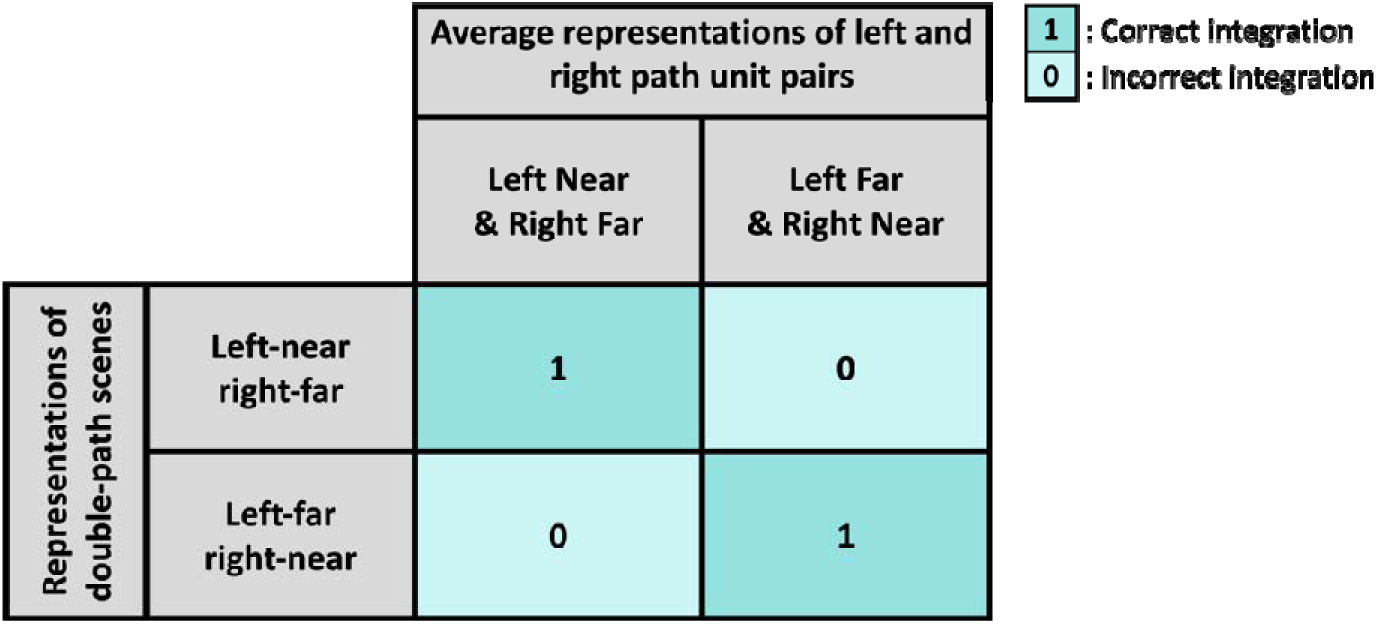
*Integrated-features* approach contrast. Contrast conditions assessing the representation of direction and distance in multiple-path scenes. Neural similarities of synthesized representations using two single-path scenes that share both path units with double-path scenes (cells marked with 1) were contrasted with neural similarities of synthesized representation from single-path scenes that do not share any path unit with double-path scenes, while sharing all independent features (cells marked with 0). It is critical that all scene pairs share the same independent features (*left*, *right*, *near*, and *far*) regardless of its condition, allowing for the examination of integration of features over pooled features.

In this contrast, it is critical that even the scene pairs in the *incorrect integration* condition share the same number of individual features with the *correct integration* condition, only not as a whole path unit. Figure 5 shows that scene pairs included in this analysis all share *left, right, near,* and *far* features, despite the pairs being separated into different conditions. Therefore, this contrast examines the exact binding of distances onto its corresponding directions, as opposed to the mere existence of such features disregarding the integration of the distance onto a specific direction. We hypothesized that if the OPA’s representation of navigational distance is precisely mapped onto its corresponding direction in a multiple-path scene, then the neural similarity will be higher for the *correct integration* condition compared to the *incorrect integration* condition. We conducted a 2 2 ANOVA to test for the interaction between the regions-of-interest (ROIs) and the conditions. We then contrasted the two conditions particularly in the OPA, and significance levels were FDR corrected.

### Integration of Path Units into a Scene: Integrated-paths approach

In the *integrated-paths* approach, we conducted an analysis using voxel-based linear combination to investigate the mechanism behind the integration of separate paths in one visual scene. The method for synthesizing the average pattern of two single-path scenes used in the *integrated-features* approach was applied here, expanding to all four single-path scene combinations and all four double-path scenes. If the OPA represents double-path scenes as a combination of the two corresponding single-path scenes, then the neural similarity will be higher between single-path scene combinations that share more path units with the double-path scenes (Baldassano et al., 2017; Kaiser, Strnad, Seidl, Kastner, & Peelen, 2014; MacEvoy & Epstein, 2011).

In the *two shared paths* condition (cells marked with 2 in Figure 6), the neural similarities between combinations of two single-path scenes that match all the paths of a double-path scene are included (e.g., the average representation of a *left near* and a *right near* single-path combination with the representation of a *left-near right-near* double-path scene). The *one shared path* condition (cells marked with 1 in Figure 6) includes neural similarities between combinations of two single-path scenes that match of the two paths of a double-path scene (e.g., average representation of a *left near* and a *right near* single-path combination with the representations of *left-near right-far* and *left-far right-near* double-path scenes). The *no shared path* condition (cells marked with 0 in Figure 6) consists of neural similarities between combinations of two single-path scenes and a double-path scene that do not match the paths of a double-path scene (e.g., average representation of a *left near* and a *right near* single-path combination with the representation of *left-far right-far* double-path scene).

**Figure 6.**
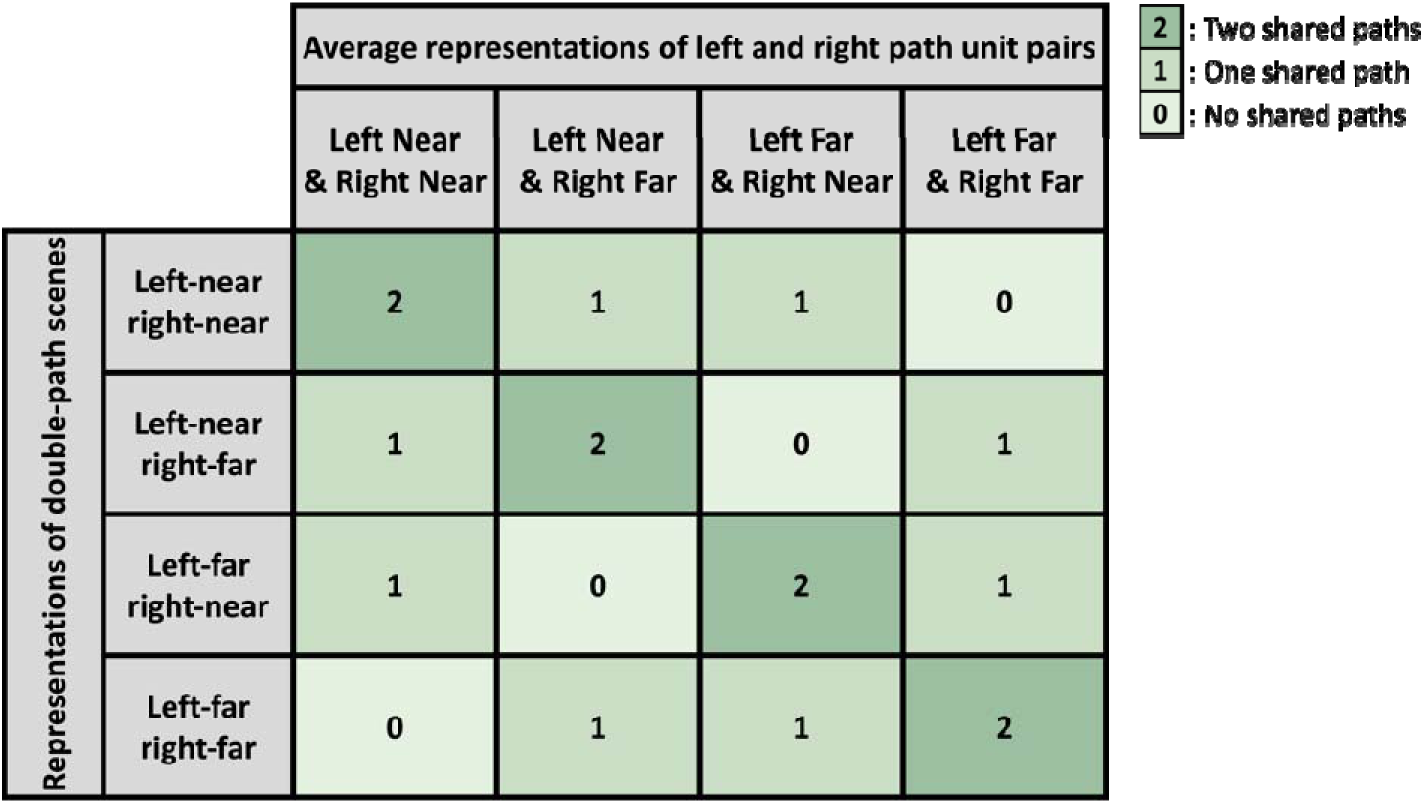
*Integrated-paths* approach contrast. Contrast conditions assessing the representation of path units within an entire scene. The *two shared paths* condition includes neural similarities of the average of two single-path scenes that combines to make the same paths as a double-path scene (same distance on the same direction; cells marked with 2). The *one shared path* condition includes neural similarities of the average of two single-path scenes that share one path unit with a double-path scene (cells marked with 1). The *no shared path* condition includes neural similarities of the average of two single-path scenes that do not share any path unit with a double-path scene (cells marked with 0).

Figure 7 depicts an example of the three conditions. The *left-near right-far* double path scene shares both paths with the *left near* and *right far* single path combination and shares one path with the with the *left near* and *right near* single path combination. It shares no path with the *left far* and *right near* single path combination.

**Figure 7.**
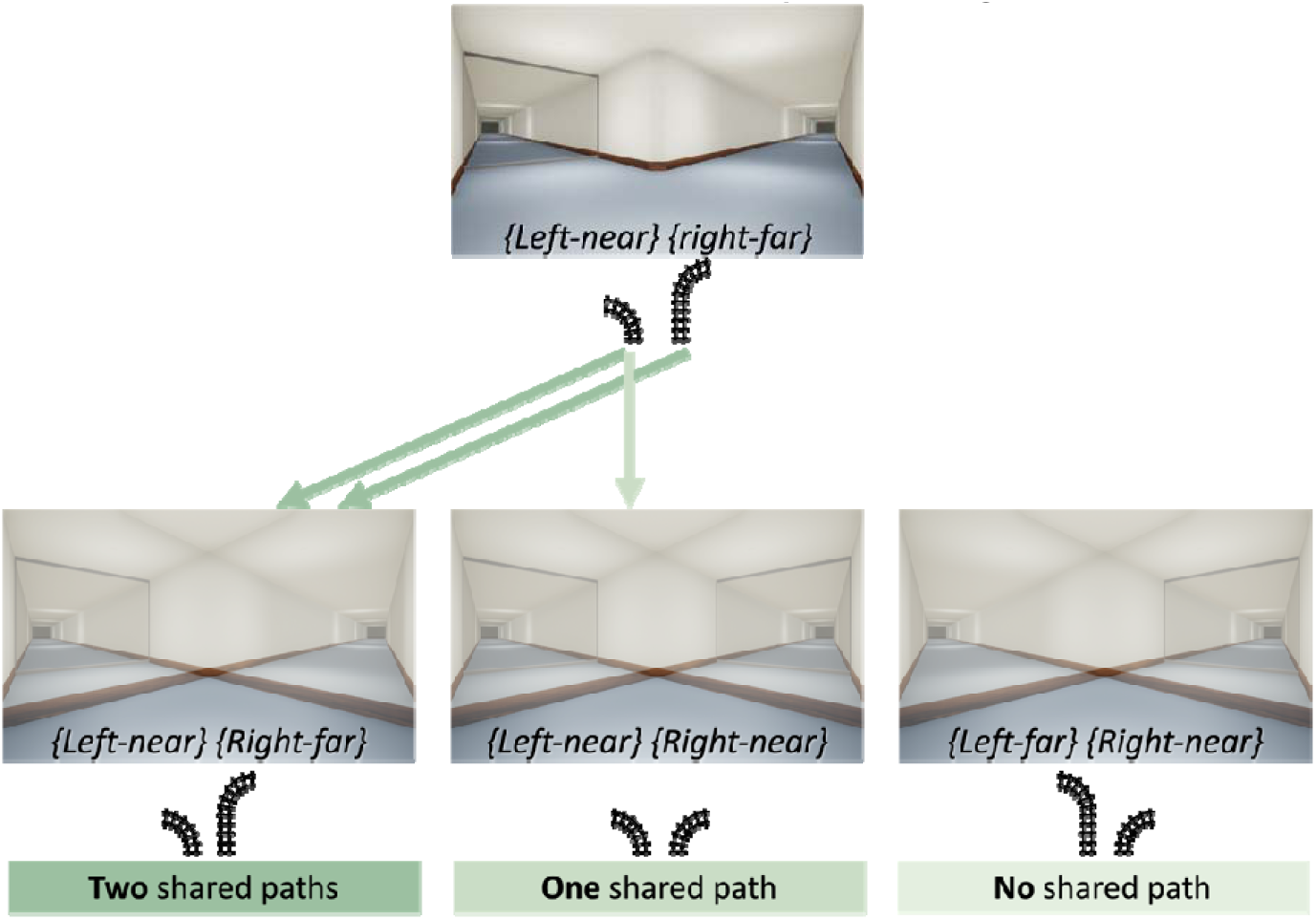
An illustration of an example for the *two shared path* condition, *one shared path* condition, and the *no shared path* condition. Each rail represents a path unit, a combined form of direction and distance features.

On average, the scene pairs in the *one shared path* condition share an equal number of each feature with the *no shared path* condition. In the *one shared path* condition, all pairs share one of *near* or *far* distance features. Two pairs in the *no shared path* condition share both and *near and far* distance features and the other two pairs share no distance features, resulting in an average of one shared feature. Therefore, if the paths in a scene are represented as the pooled sum of each feature, we expect no significant difference between these two conditions. An performed a 2 3 ANOVA to test if the response pattern of the OPA is different from the response pattern of the PPA. We then contrasted the three conditions particularly in the OPA, using FDR to correct for the significance levels.

## METHODS

### Participants

Twenty-four participants were recruited from the Yonsei University community with financial compensation (19 female, *M* = 24.54, *SD* = 2.65). The sample size was estimated with a medium effect size (partial of 0.06) based on past fMRI studies investigating into scene selective regions. Two participants were excluded from the analysis because their ROIs were not localized. Runs with excessive head motion (motion correction parameter exceeding 3mm within a run) were excluded from the analysis (four out of ten runs from one participant; two out of ten runs from another). Another participant was excluded from the analysis due to low task performance that indicates absence of basic attention to the presented stimuli (number of missed trials in a detection task exceeding 2 standard deviations from the mean). All analyses were carried out with data from 21 participants. All participants had normal or corrected-to-normal vision. Written consent was obtained from all participants, and study was approved by the Yonsei University Institutional Review Board (IRB).

### Stimuli

Artificial indoor scenes with corridors were created and captured in first-person view using Unreal Engine 4. There were eight different stimulus conditions (2: number of paths

2: directions 2: distances). There were 24 variations of wall and floor textures within each type of scenes. In single-path scenes, the path was directed either towards the left or towards the right (Figure 8a). In double-path scenes, left and right paths were both present while a viewer stood at a crossroad. (Figure 8b). Each path contained a glass wall that was located either near or far, changing the navigable distance of each path. All stimuli were displayed in 1152 648 pixels (21.2° 12.0° visual angle).

**Figure 8.**
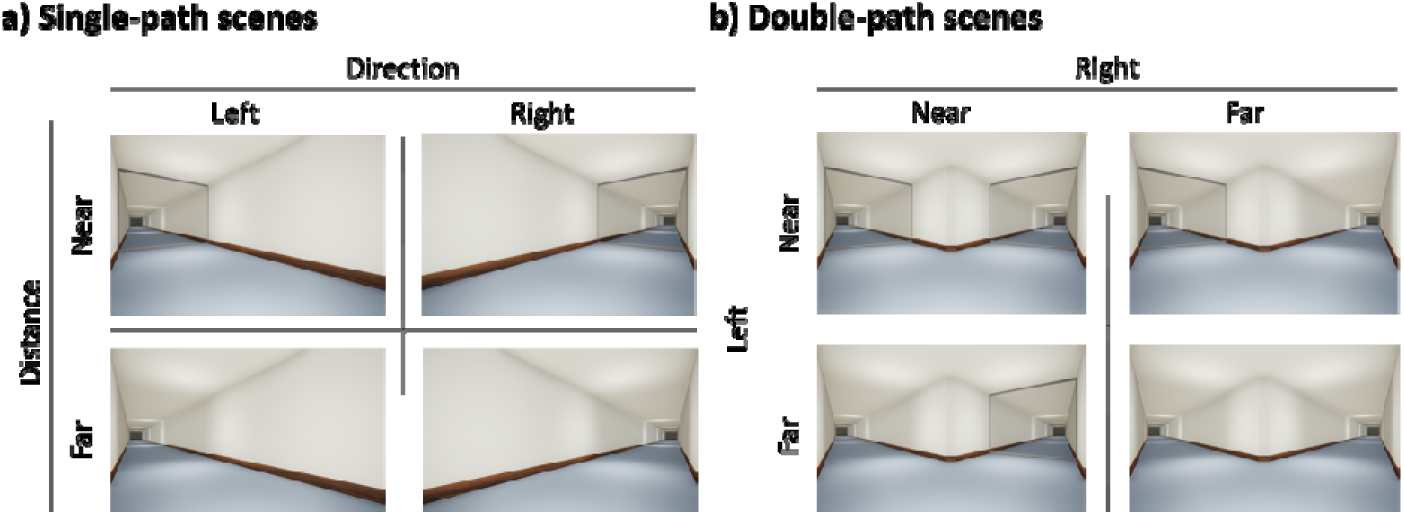
Examples of images in the eight stimulus conditions. (a) Four single-path scenes, each with a straight corridor directed towards the left or towards the right, and a glass wall located either near or far. (b) Four double-path scenes, each with two corridors, one directed towards the left and the other directed towards the right. A glass wall is located either near or far in each path.

### Procedure

The experiment consisted of 10 experimental runs (5.47 minutes, 164 repetition time (TR) each). An experimental run consisted of 16 blocks presented in a random order (two blocks per stimulus condition). In each block, 12 images from one of the eight stimulus conditions were presented serially. Each image was displayed for 600ms followed by a 400ms interstimulus interval (ISI) with a fixation dot. The image sequence within each block was randomized. The stimulus condition blocks were interleaved by an 8-second fixation period. While viewing the stream of images, participants performed a frame detection task where they pressed a button when an image was presented in a red frame (Figure 9). The red frame task was used instead of conventional fixation detection task or one back repetition task to keep the participants’ attention to the whole view of a scene (Park, Josephs, & Konkle, 2022; Tarhan & Konkle, 2020). Participants were instructed to pay attention to the navigable directions and distances of paths in the presented scene images while performing the frame detection task.

**Figure 9.**
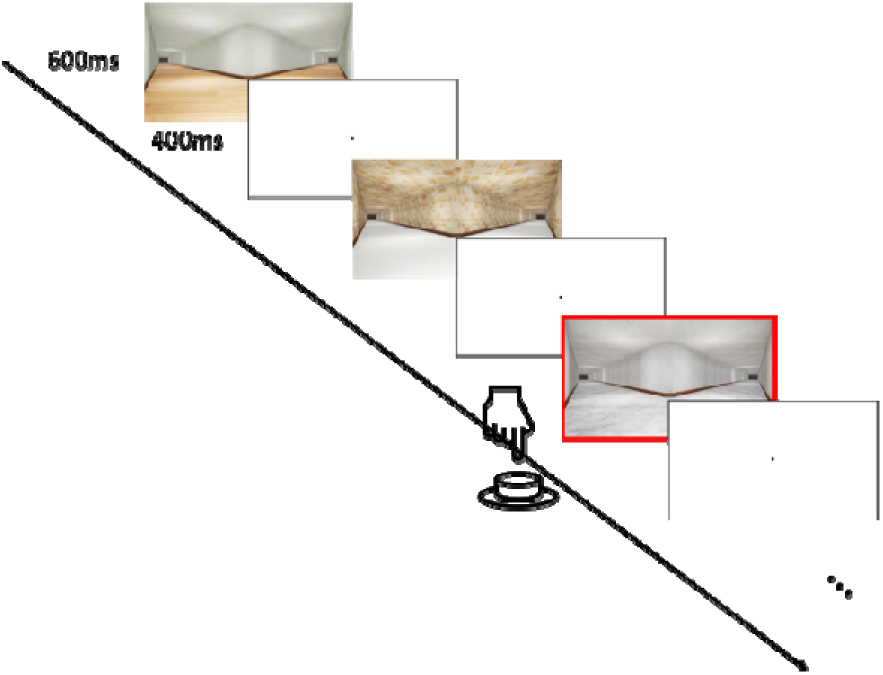
The schematic visualization of the procedure within a condition block. A series of 12 images were presented from one of the eight conditions. Participants pressed a button when a red frame was presented along the border of the image, while paying attention to the path direction and navigational distance. There were 16 condition blocks in a run, and each participant went through ten experimental runs.

### MRI Acquisition

The fMRI data was acquired using Philips Ingenia 3.0T CX scanner with a 32-channel head-coil, located at the Yonsei University Convergence Medical Technology Center. Structural T1-weighted images were acquired by magnetization-prepared rapid-acquisition gradient echo (MPRAGE) with 0.5 X 0.5 X 0.5 mm voxels. Functional image was acquired through a gradient echo T2* weighted sequence (2.625 X 2.625 X 3 mm voxels; TR = 2 s; TE = 22 ms; flip angle = 90°; 42 axial slices), acquired parallel to the anterior commissure-posterior commissure (ACPC) line. During the scan, a wedge leg support cushion was placed under the knees of the participants to reduce the transfer of leg motion to the head.

### Preprocessing

Brain Voyager software (Brain Innovation, Maastricht, Netherlands) was used for preprocessing of the functional data. The preprocessing included slice time correction, motion correction, and temporal high-pass filtering including linear trend removal. Spatial smoothing was not included. The data was aligned to the Montreal Neurological Institute (MNI) space after co-registering it to the structural image.

### Regions of Interest

Individual OPAs were identified using a functional localizer run (each 7 minutes 6 seconds, 213 repetition time (TR)) independent from the experimental runs (Kamps, Lall, & Dilks, 2016; Pitcher, Dilks, Saxe, Triantafyllou, & Kanwisher, 2011). The localizer run started and ended with 10-second fixation blocks and took place after the fifth experimental run in each participant. 3-second video clips of faces, scenes, objects, and scrambled objects (720 X 480 pixels) were presented in an fMRI block design. A total of 16 stimulus blocks were presented, with four blocks for each of the four stimulus conditions. Latin square design was used to determine the order of the stimulus blocks. Eight videos were presented in each stimulus block. Each video was displayed for 3000 ms followed by 500 ms of blank screen. 10-second fixation blocks were interleaved between every four stimulus blocks. No task that required responses from the participants was performed, but they were instructed to pay attention to the videos.

The OPA and the PPA were defined by contrasting the level of brain activity during scene blocks to activity during face blocks (Scenes > Faces) from the dynamic localizer run (*p* < .0001, uncorrected, Epstein & Kanwisher, 1998). Bilateral ROIs were identified by selecting voxel clusters that passed the threshold near the transverse occipital sulcus for the OPA, and the posterior parahippocampal gyrus for the PPA. We were able to identify bilateral OPA from all 21 participants included in the analysis, except for the right OPA of one participant. Bilateral PPA was also identified from all 21 participants, except for the right PPA of one participant, and the left PPA of another participant.

We also examined the EVC as a control region for low level visual features. The EVC was defined by contrasting the level of activity during scrambled object blocks to activity during intact object blocks (Scrambled objects > Objects) from the same functional localizer used in identifying the OPA and the PPA (*p* < .0001, uncorrected, Coggan et al., 2017; Bonner & Epstein, 2017). We identified bilateral EVC from eight participants, and one of the left or the right EVC from another seven participants. No interactions were observed between contrast conditions and the left and the right ROIs, so they were collapsed for all the analyses.

### Neural Similarity Analysis

The MRI signal intensity values were extracted and normalized within runs in each ROI, allowing the analysis across runs. In ROIs of each subject, the activities of each voxel were averaged across the TRs within the same stimulus condition block, after adjusting for the hemodynamic lag of 4 seconds (2TR). This resulted in generating a multi-voxel pattern for each stimulus condition in each ROI of each subject.

This pattern was analyzed using the MATLAB toolbox for RSA (Kriegeskorte, Mur, & Bandettini, 2008; Nili et al., 2014). An 8 X 8 representational dissimilarity matrix (RDM) was generated, where each element of the matrix indicated the representational distance (1 - Pearson correlation coefficient) between the patterns for each stimulus condition. We applied predefined contrasts from the *independent-features-*, *integrated-features-,* and the *integrated-paths* approaches to perform hypothesis testing across conditions.

## RESULTS

### Independent Representations for Direction and Distance: Independent-features approach

In the *independent-features* approach, we examined whether direction and distance of a path are represented simultaneously in the OPA (Figure 1). To do this, we conducted a 2 X 3 ANOVA contrasting the neural similarities of single-path scene pairs that had the same direction, pairs with the same distance, and pairs that did not share any feature in the OPA and the PPA. The result revealed a marginally significant interaction between conditions and ROIs (*F*(2,40) = 2.868, *p* = .030). More importantly, this analysis revealed that the OPA’s multi-voxel response codes information about the path direction and distance (Figure 10). Specifically, there was a significant difference between the *shared direction* condition and the *no shared feature* condition in the OPA (*t*(20) = -2.668, *q* = .015). For example, the representation for a *left near* path is more similar to a *left far* path compared to the representation of a *right far* path. There was also a significant difference between the *shared distance* condition and the *no shared feature* condition in the OPA (*t*(20) = -2.392, *q* = .041). For example, the representation for a *left near* path is more similar to a *right near* path compared to the representation of a *right far* path. There was no significant difference between the *shared direction* condition and the *shared distance* condition (*t*(20) = -0.105, *q* = .918). Overall, the results suggest that path direction and navigational distance are both coded for single-path scenes in the OPA, replicating the results of Epstein and Bonner (2017) and Park and Park (2020). Furthermore, the OPA simultaneously takes both direction and distance features into account to a similar degree in representing a visual path.

**Figure 10.**
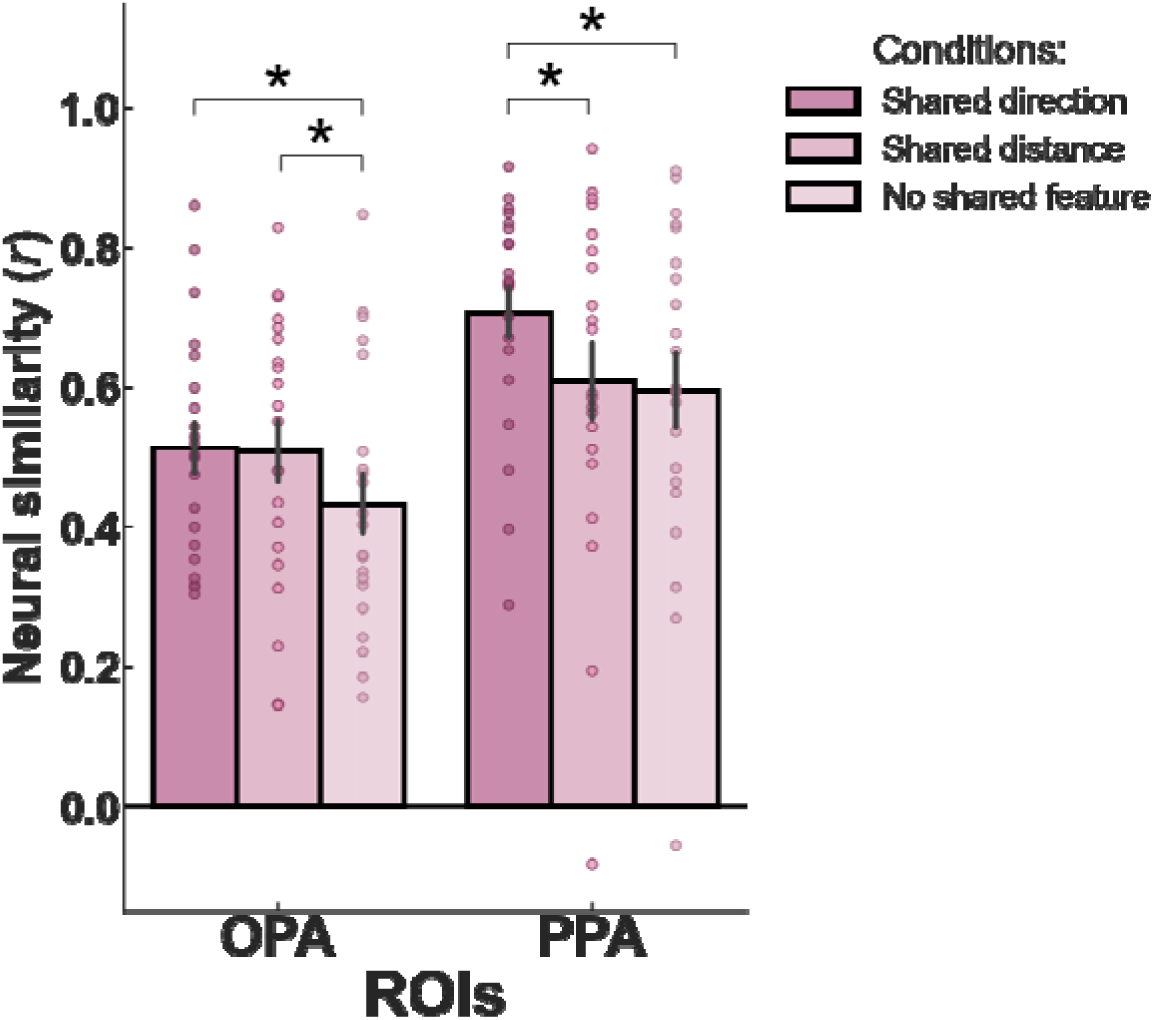
Results for the *independent-features* approach contrasts. Analysis comparing the neural similarity of the *shared direction* condition, the *shared distance* condition, and the *no shared feature* condition. When comparing with the *no shared feature* condition, similarity in the *shared direction* condition and the *shared distance* condition were both significantly higher. A direct comparison of the *shared direction* and *shared distance* conditions in the OPA was not significant, showing no evidence for the bias for one feature over another. \**q* < 0.05

There was a dissimilar pattern between the three conditions in the PPA, suggesting the unique function of OPA in representing features related to navigation. An analysis within the PPA showed a significant difference between the *shared direction* and the *no shared feature* conditions (*t*(20) = -3.731, *q* < .006). There was no significant difference between the *shared distance* condition and the *no shared feature* condition (*t*(20) = -0.730, *q* = .474). The *shared direction* condition and the *shared distance* condition were significantly different (*t*(20)= -3.503, *q* < .006). From this result and a previous finding that the PPA is not sensitive to path direction (Dilks et al., 2011), it can be implied that the PPA reflects place similarity, considering that a glass wall can be installed to or removed from the same place, but different configurations of walls indicate different places.

Taking the findings from past research into consideration, the differences between conditions in the OPA likely reflect functional differences for visual navigation more so than the low-level features of the images (Dilks et al., 2011; Epstein & Bonner, 2017; Park & Park, 2020; Persichetti & Dilks, 2016). To account for the effect of low-level features in the response pattern of the OPA, we created a low-level image similarity model using pixel-wise root mean square difference (RMSD) for all single-path scene pairs (Erez et al., 2016). The model was compared with the responses of OPA and EVC of each participant using Kendall’s rank correlation. We then performed a paired *t*-test to examine whether the OPA and EVC had significantly different correlations with the RMSD model (Figure 11). If the OPA’s response pattern is beyond mere reflection of low-level image similarities, then the OPA is expected to have a lower correlation with the RMSD model compared to the EVC. The result showed that the RMSD model had a significantly lower rank correlation with the OPA’s similarity pattern compared to the rank correlation with the EVC’s response pattern (*t*(14) = -2.630, *p* = .020). This suggests that the OPA’s response pattern for navigationally relevant features are not driven solely by the low-level visual similarities of the images.

**Figure 11.**
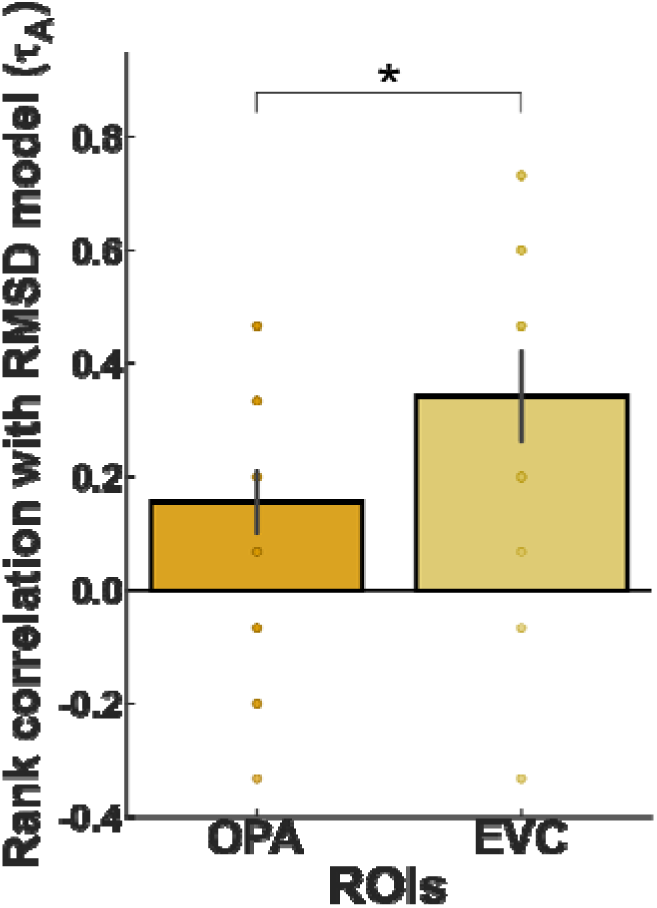
Rank correlation of OPA and EVC with the RMSD model. The correlation between the model and the OPA’s response pattern was lower than the EVC. \**p* < 0.05

### Combined Representation of Directions and Distances: Integrated-features approach

In the *integrated-features* approach, we conducted a 2 2 ANOVA to compare the neural similarities between the average representations of two single-path scenes of different directions and distances to the representations of double-path scenes with different distances on each path (Figure 5). The result showed a significant interaction between the conditions and the ROIs (*F*(1, 20) = 7.410, *p* = .013). Most critically, the OPA’s representations of distances were bound to its corresponding directions, beyond the coding of mere existence of these features (Figure 12). Specifically, the neural similarities between the representations of synthesized average and the double-path scenes in the *correct integration* condition were higher than those in the *incorrect integration* condition in the OPA (*t*(20) = -2.904, *q* = .009). For example, the representation of a *left-near right-far* double-path scene was significantly more similar to the representation of a *left near* or a *right far* single-path scene compared to the representation of a *left far* or a *right near* single-path scene. This result shows that the OPA’s representation of each of the two paths in double-path scenes is related to the representation of single-paths with the same distance on the same direction, as units.

**Figure 12.**
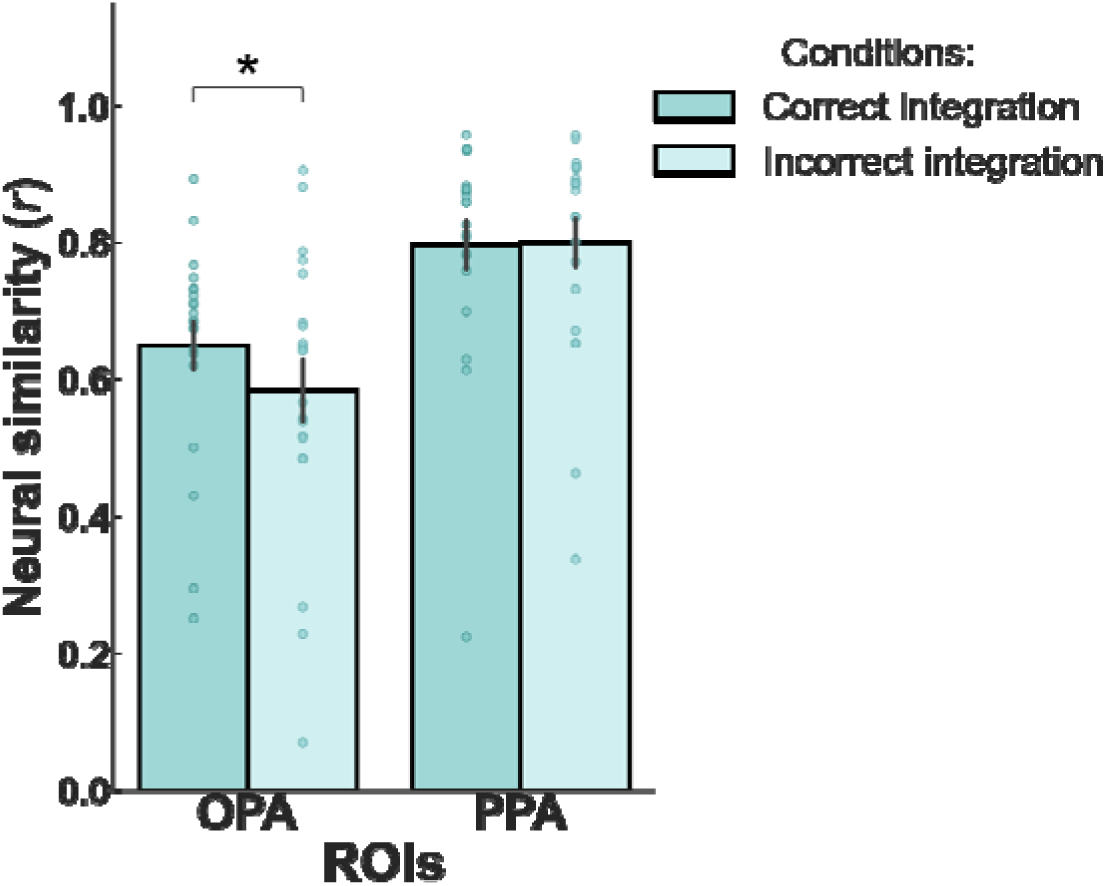
Results for the *integrated-features* approach contrast. Analysis comparing the neural similarities of an average of two single-path scenes and a double-path scene pairs in the *correct integration* and the *incorrect integration* conditions. The difference between conditions was significant only in the OPA. \**q* < .05

To confirm that the integration of direction and distance features are unique to the OPA, we compared the *correct integration* and the *incorrect integration* conditions in the PPA, another scene-selective region. Unlike the OPA, there was no significant difference between the *correct integration* and the *incorrect integration* conditions in the PPA (*t*(20) = 0.167, *q* = .869). This shows that the integration of direction and distance features into a unit is not an inherent property of scene-selective areas in general but is rather unique to the function of the OPA.

### Integration of Path Units into a Scene: Integrated-paths approach

In the *integrated-paths* approach, we further investigated how the OPA integrates multiple paths within a visual scene (Figure 6). We compared the neural similarities between the average representation of two single-path scenes of different directions to representation of a double-path scenes in the *two shared path*, *one shared path*, and the *no shared path* conditions in the two ROIs using a 2 X 3 ANOVA (Figure 13). After correcting for the violation of sphericity (Mauchly’s test, x^2^(2) = 33.870, *p* < .001) using the Greenhouse-Geisser estimates of sphericity (E = .546; Greenhouse & Geisser, 1959), the result revealed a significant interaction between the three conditions and the two ROIs, (*F*(2.73, 54.6) = 9.791, *p* < .001). Most critically, the result revealed significant differences across conditions particularly in the OPA. The OPA’s average representation of two single-path scenes was significantly more similar to a double-path scene in the *two shared paths* condition compared to the *one shared path* condition (*t*(20) = -2.841, *q* = .010) and compared to the *no shared path* condition (*t*(20) = 3.521, *q* = .010). The similarity was also higher for *one shared path* condition compared to the *no shared path* condition (t(20) = 3.82, *q* = .010).

**Figure 13.**
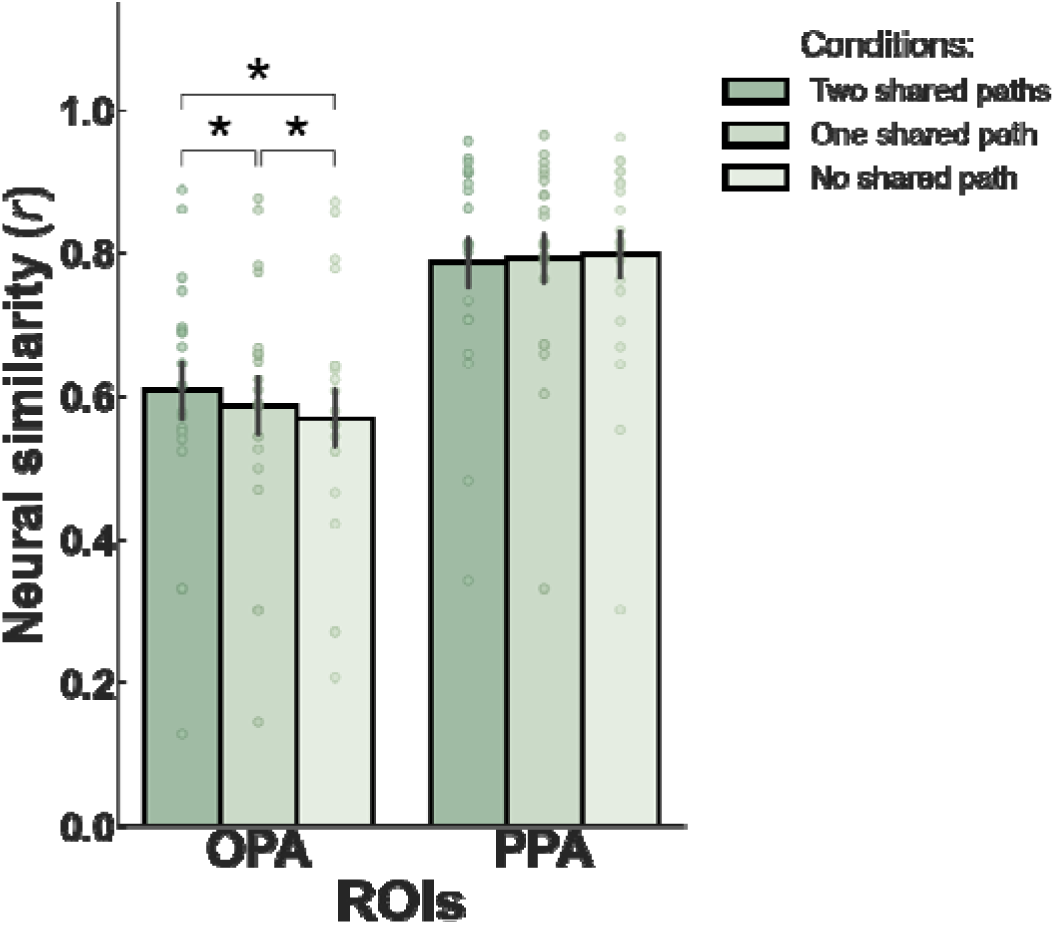
Results of the *integrated-paths* approach contrast. Analysis comparing similarities between the average representation of two single-path scenes and a double-path scene in the *two shared path* condition, *one shared path* condition, and *no shared path* condition. The OPA showed the highest neural similarity for the *two shared path* condition, and the lowest for the *no shared path* condition, with the *one shared path* condition in between. All differences among the three conditions were significant in the OPA. \**q* < 0.05

Note that the scene pairs in the *no shared path* condition share the equal amount of individual features on average compared to the *one shared path* condition, once again confirming that the OPA represents paths as a combined unit of direction and distance, rather than as the sum of all individual features. Overall, the results suggest that the representations of each path in the double-path scenes are based on the combination of two specific single-path units. Specifically, the representation of a double-path scene (e.g., *left near right-far* scene) is built based on the combination of each path unit, with specific distance and direction features combined (e.g., {*left near*} & {*right far*}) rather than a pooled representation of all features (*left* & *right* & *near* & *far*).

To test whether this pattern of result is found generally for any scene-selective area, we compared the three conditions in the PPA. For the PPA, there was no significant difference between the *two shared path* condition and the *one share path* condition (*t*(20) = -0.884, *q* = .378). There was also no significant difference between the *two shared path* and the *no shared path* conditions (*t*(20) = -1.101, *q* = .341), and between the *one shared path* condition and the *no shared path* condition (*t*(20) = -1.207, *q* = .341). This again suggests that combining the representation of each path into a scene is not a general function found across scene-selective areas.

## DISCUSSION

In this study, we aimed to investigate how the OPA represents paths in a visual scene. First, in the *independent-features* approach, we confirmed that directions and distances of navigable paths are simultaneously represented by the OPA, with no significant bias for one feature over another. In the *integrated-features* approach, we showed for the first time that the OPA correctly puts together a direction feature and a distance feature into a path unit. Last, we expanded the scope of our analyses by using the neural similarities of all possible pairs of scenes in the *integrated-paths* approach. Here, we demonstrated that the representations of scenes with multiple paths are formed based on the combination of each path unit within that scene, and confirmed once again that direction and distance features are correctly bound by the OPA.

This correct binding of directions and distances is a requisite for the execution of visually guided navigation. Inability to bind these features correctly may result in the failure to calculate whether and how far one can move towards a specific direction. The results of our experiment suggest that the OPA supports the recognition of available path trajectories within the immediate visual scene by combining independent visual features that are relevant for navigation. This provides further evidence that the OPA supports visually guided navigation by rapidly computing individual features and putting together navigation related features to build a *navigation file*.

Previous work regarding neural mechanisms behind scene perception and visually guided navigation have focused more on identifying individual features that support these functions (Cheng, Walther, Park, & Dilks 2021; Henriksson, Mur, & Kriegeskorte, 2019; Persichetti & Dilks, 2016; Walther & Shen, 2014). There are some research that explored how multiple features within a scene are represented in the brain. For example, Henriksson et al. (2019) have directly compared the effects of two independent features, layout and texture, in the OPA and the PPA. They demonstrated that the OPA is more sensitive to changes in scene layout than it is to changes in texture. Although this research took multiple features into account, its main purpose was to contrast which feature provides better discriminability in the ROIs rather than to show how multiple features are simultaneously coded by an area.

Another work has shown evidence supporting the object-based channel for scene category recognition in the lateral occipital (LO) cortex, widely known for object selectivity (MacEvoy & Epstein, 2011). The response pattern of the LO for a complete scene image was accurately predicted by the linear combination of response patterns for multiple object images from the scene context. For example, the LO’s representation of a kitchen was better predicted using the linear average pattern of a refrigerator and a stove than when using the average pattern of a traffic light and a car. This research, however, has focused more on how object information, as a complete unit of features, is combined across an entire scene rather than the combination of features that reside within a complete entity. Here, we show for the first time that the OPA not only codes geometrical features of visual scenes that are especially related to navigation, but also correctly combines them to represent complete forms of paths within a scene.

Our method of combining two single-path scenes to compare with double-path scenes entails several implications for how the OPA integrates two independent features. First, the integration of navigationally related features in the OPA may be an automatic process. Putting together multiple features to build a bigger unit may be an effective computational mechanism that supports effortless navigation when there are multiple features that can be incorrectly integrated. However, the process of integration is rather unnecessary for single-path scenes, considering that they hold one direction feature and one distance feature, creating an unambiguous situation for navigation even at a pooled feature state. Nevertheless, the OPA had an integrated coding of features in both single-path and double-path scenes, shown by a higher the neural similarity between single-path scenes and double-path scenes with more shared path units over scenes with less path units. The automatic processing of OPA has been suggested previously, by showing that navigational information is represented quickly and in the absence of a navigation related task (Bonner & Epstein, 2017; Henriksson et al., 2019; Suzuki, Kamps, Dilks, & Treadway, 2021). Our work may add to these previous findings and suggest that even features in unambiguous, single-path, scenes are automatically integrated by the OPA .

Second, our method reveals an effective shared coding scheme for paths across noticeably different places of single-path scenes and multiple-path scenes rather than remapping after changes in places. This suggests an abstract processing of the OPA by using features that are common in scenes. This summarization of complex scenes resembles the spatial envelope model for scene recognition (Oliva & Torralba, 2001). The spatial envelope model proposes an abstract descriptor for a scene’s shape, treating it like an object with a unitary shape. A spatial envelope is derived from properties such as naturalness, openness, and expansion. These features together build a low-dimensional representation of natural scenes, called scene *gist*, which provides enough information for people to recognize its semantic category even at a glance (Oliva & Torralba, 2006). Like the spatial envelope model accounting for scene category recognition, the *navigation file* might account for the process of visually guided navigation. Taken together, the OPA may automatically summarize scene properties that efficiently generalizes across different places.

Keeping in mind that we used a highly controlled stimuli set, results of the current experiment may entail a low generalizability to the dynamically changing view during active navigation. Accordingly, it is important not to make a hasty conclusion that the representation of path units directly leads to navigational behavior, considering that multiple regions involved in scene processing form networks that serve different functions like navigation and putting fragments of scenes into a broader spatial context (Baldassano, Esteva, Fei-Fei, & Beck, 2016). Future work may elucidate how the combined representation of visual features in the OPA operate within a network to support active and dynamic visual navigation across the immediate and a broader environment.

## Conclusion

In conclusion, we have demonstrated that the OPA has a conjoined representation of two independent features relevant for navigation, direction and distance, for complex double-path scenes as well as simple single-path scenes. This type of coding was not evident in the PPA, a different scene-selective region, and its process was beyond a simple reflection of low-level visual features. The conjunction of features in the OPA seems to be automatic, and this conjunctive coding scheme of navigational features is generalizable across different places, likely supporting efficient visually guided navigation.

## Acknowledgements

This research was supported by the National Eye Institute (R01EY026042); and the National Research Foundation of Korea (NRF-2023R1A2C1006673) to SP.

